# *In vitro* generation of human embryonic stem cell-derived heart organoids possessing physiological ion currents

**DOI:** 10.1101/2022.05.15.491904

**Authors:** Jiyoung Lee, Hiroshi Matsukawa, Kohei Sawada, Rin Kaneko, Fumitoshi Ishino

## Abstract

Recently, methods for *in vitro* organogenesis have been broadly developed due to their strong potential for human applications in medicine. In the present study, we optimized the culturing method of human heart organoids (hHOs) from embryonic stem cells (ESCs) in the presence of the highly concentrated laminin-entactin and fibroblast growth factor 4. The resulting hHOs showed distinctive cardiac morphology with atrium- and ventricle-like chambers composed of cardiac cells as well as expressed the integral proteins of gap junctions and ion channels. In fact, isolated cardiomyocytes from these hHOs exhibited Na and Ca currents by patch clamp analysis. These results indicated that the present method will provide a powerful tool for cardiac safety assessment of newly developed drugs as an *in vitro* human ESC-derived test system.

**One-Sentence Summary:** FGF4 and ECM contribute to the generation of human HOs with heart compartments and electrophysiological properties.

## Main Text

Heart organoid-generating systems have potential applications in drug screening, regenerative medicine and structural and functional analyses of heart defects. Various differentiation protocols of cardiac cells have been developed: 1) *in vitro* cardiomyocyte differentiation from pluripotent stem cells [induced pluripotent stem cells (iPSCs) and embryonic stem cells (ESCs)](*1–4*) and 2) *in vivo* and *in vitro* direct reprogramming of cardiomyocyte-like cells from fibroblasts (*5–7*). Given the increasing demand for the generation of structurally organized and functional heart organoids, we recently developed a method for three-dimensional (3D) heart organoids (*8*) with *in vivo* heart-like structures as well as cardiac functions implying electrophysiological properties by culturing mouse embryoid bodies (EBs) from ESCs with the laminin-entactin (LN/ET) complex (*9*) and fibroblast growth factor 4 (FGF4) (*10*). Translational studies of the mouse heart organoid-generating system are necessary for human applications, and we established a method for human PSC-derived heart organoids based on the previous laminin and FGF4 method. Using human ESCs was a reasonable choice for easier maintenance of the undifferentiated state than human iPSCs; nevertheless, hESCs have been under ethical issues. In this study, we improved the heart organoid culture conditions and generated hESC-derived heart organoids with cardiac structural features and an electrophysiological capacity for heart function. The generated hESC-derived heart organoids possess gap junction proteins required for the intercellular conduction system in the heart as well as critical ion channel proteins for generating action potentials, including K_ir_ 2.1, hERG, Na_v_ 1.5 and Ca_v_ 1.2. Furthermore, the observation of functional Na and Ca ion currents in patch clamp methods indicates the feasibility of the human heart organoids generated by the present method can be used for nonclinical cardiotoxicity screening.

## Results

### Heart organoid generation from human embryonic stem cells (ESCs)

In a previous study, we generated functional 3D heart organoids from mouse ESCs in the presence of the laminin-entactin (LN/ET) complex via the exogenous growth factor, fibroblast growth factor 4 (FGF4) *in vitro*. These mouse heart organoids exhibited functional activities, including beating movement, cardiac muscle contraction and electrophysiological characteristics such as action potential propagation. Given that *in vitro* mouse heart organoids showed appropriate drug responses in electrophysiological studies, these organoids will be useful for preclinical cardiotoxicity screening. However, because of the different composition of ion channels and the dynamics of intracellular Ca as well as the different heart rate between rodents and human cardiac muscle (*11*), it is necessary to use biomimetic materials that reproduce human cardiac tissue instead of rodents, in order to overcome the problem of species difference (especially rodents) in the field of drug discovery. Particularly, the use of human ES derived heart organoids as surrogates for human cardiomyocytes (CMs) as well as human hearts would be more appropriate methodology for the evaluation of cardiac toxicity in newly developed drug screening.

Therefore, we generated heart organoids by using human ESCs (hESCs). The hESCs were cultured and maintained with bFGF (10 ng/mL) in primate ESC culture medium, and then, embryoid bodies (EBs) were formed without bFGF in KSR-EB medium. To determine whether the LN/ET complex and FGF4 signalling are also key factors for the self-organization of human heart development, we cultured intact hESC-derived EBs on gelated LN/ET complexes with FGF4. Given that laminin 111 of the LN/ET complex is abundantly expressed during early heart development and laminin 411 is highly expressed in mouse embryonic day 12.5 (E12.5) hearts (*8, 12*) (fig. S1A), we exchanged extracellular matrix from the LN/ET complex with laminin 411 (iMatrix-411) at culturing day 11. As expected, the hEBs showed cardiac morphogenic changes, as shown in the mEB-derived heart organoids. Consequently, hESC-derived heart organoids with multiple chambers were efficiently generated in the human heart organoid-generating system via a two-step extracellular environment (Fig. 1A and fig. S1B)

**Fig. 1.**
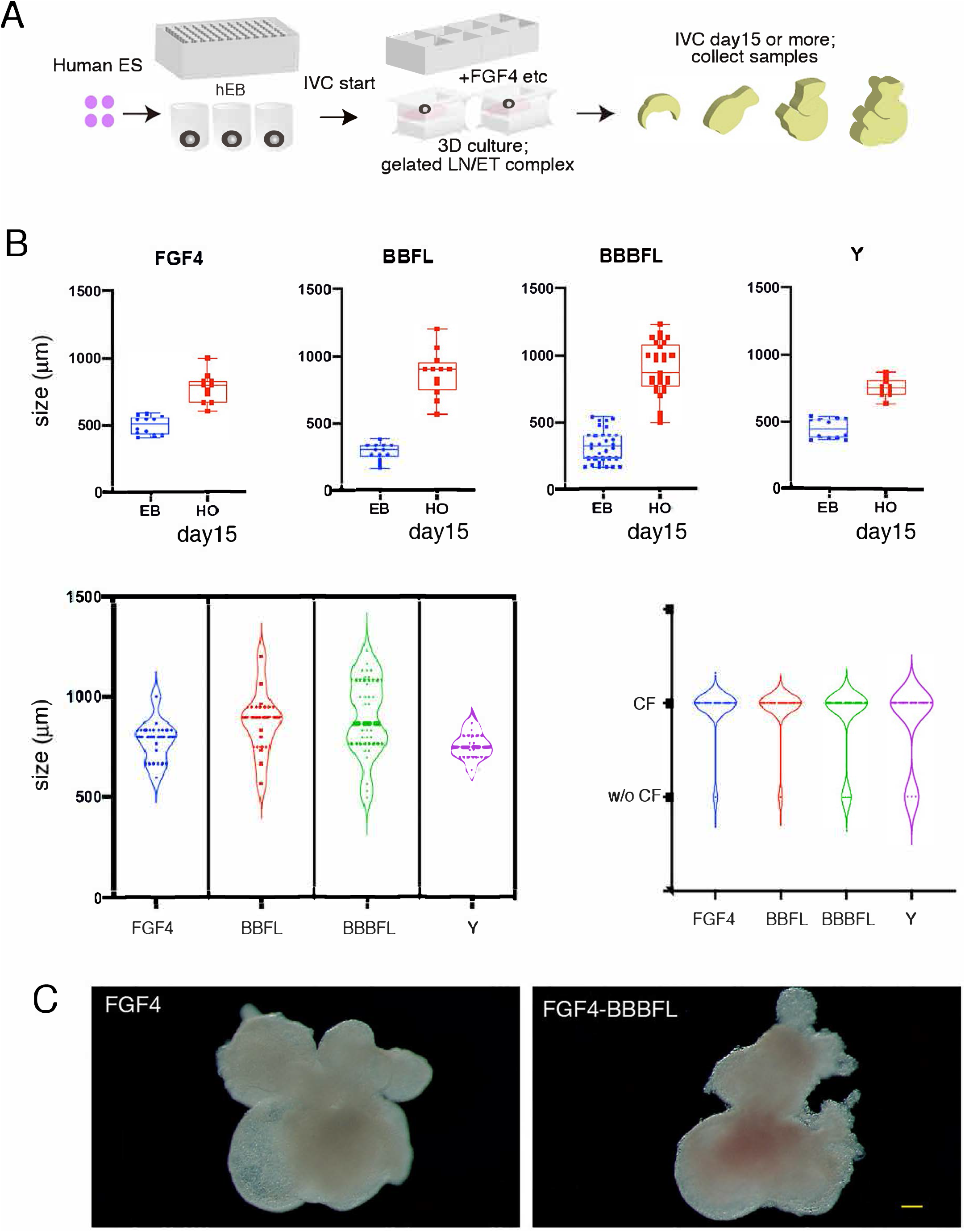
*In vitro* recapitulation of developing hearts. (**A**) Experimental scheme to generate hESC-derived heart organoids (HOs) *in vitro*. The hEBs derived from hESCs were cultured on the gelated LN/ET complex with HO medium containing FGF4. (**B**) Effect of BMP10 for HO formation. The hEBs were cultured with FGF4 only, BBFL and BBBFL conditions on the LN/ET complex (over two experiments). Chamber formation was observed under the above conditions. (**C**) Representative hESC-derived HOs with multiple heart chambers generated using the above culture system after culturing for 15 days. The left panel indicates hESC-derived HOs in the presence of FGF4 only, and the right panel indicates hESC-derived HOs in the presence of BBBFL. Both hESC-derived HOs showed remarkable morphogenic changes. See also fig. S1B. Scale bar: 100 µm.

To produce heart organoids with perfect structural similarity to the *in vivo* heart, we investigated several culture conditions, for example, FGF4 only; additional supplementation with 6’ bromoindirubin 3’ oxime (BIO, a Wnt activator), bone morphogenetic protein 4 (BMP4), and LIF (BBFL) from day 9 to day 15; BIO, BMP4, BMP10, and LIF (BBBFL) from day 9 to day 15; and additional supplementation with the ROCK inhibitor Y27632 (Y) (Fig. 1B). Because BMP10 is expressed in E9.5 and E11.5 mouse embryonic hearts despite loss of expression in previous mouse heart organoids, BMP10 is believed to be a critical factor for adequate heart development. The hEBs cultured under all conditions showed growth and morphological changes. In particular, the morphology of heart organoids at day 11 of culture from hEBs matched that of the mouse embryonic heart composed of atria and ventricles, or human fetal heart 29 days of gestation. These conditions seem to be specialized to promote self-organization for cardiac morphogenesis during the culture of heart organoids. In addition, the cultured heart organoids in the presence of BBFL and BBBFL showed vigorous proliferation compared to those with FGF4 only, suggesting that BIO, BMP4, BMP10, and LIF promote growth in FGF4-mediated heart development in this culture system. The comparison of the morphology of heart organoids between the FGF4 and FGF4-BBBFL conditions showed that BBBFL is efficient for the generation of more mature and differentiated heart organoids (Fig. 1C). Regarding the acquisition of cardiac functionality, these hESC-derived heart organoids by FGF4 and FGF4-BBBFL conditions showed beating movements (movies S1 and S2).

### Evaluation of hESC-derived heart organoids by histological analysis and ultrastructural analysis

Next, we performed histological evaluation of the generated hESC-derived heart organoids. Major sarcomere proteins in CMs, atrial myosin light chain 2 (Mlc2a) and ventricular myosin light chain 2 (Mlc2v) (*13*), were expressed in 15-day cultured hESC-derived heart organoids (Fig. 2A), although the signal of Mlc2v was weak in these heart organoids, similar to that in E10.5 mouse hearts. In addition to CMs, the heart consists of other types of cells, smooth muscle cells (SMCs) and endothelial cells (ECs), which are important for angiogenesis (blood vessel formation) in cardiogenesis. Therefore, we investigated the expression of a CM marker, cardiac troponin T (cTnT) (*14*); an EC marker, platelet/endothelial adhesion molecule 1 (PECAM; CD31) (*15*); and an SMC marker, 𝔞-smooth muscle actin (αSMA), to determine whether the organization of cardiac structural cells occurs properly in hESC-derived heart organoids. As a result, whole-mount immunostaining using an optimized clear, unobstructed brain/body imaging cocktail and computational analysis (CUBIC) method (*16*) showed the embryonic heart-like organization of cardiac structural cells in the inner structure of the hESC-derived heart organoids (fig. S2), suggesting that this culture system enables the *in vitro* generation of heart organoids from hESCs *in vitro*.

**Fig. 2.**
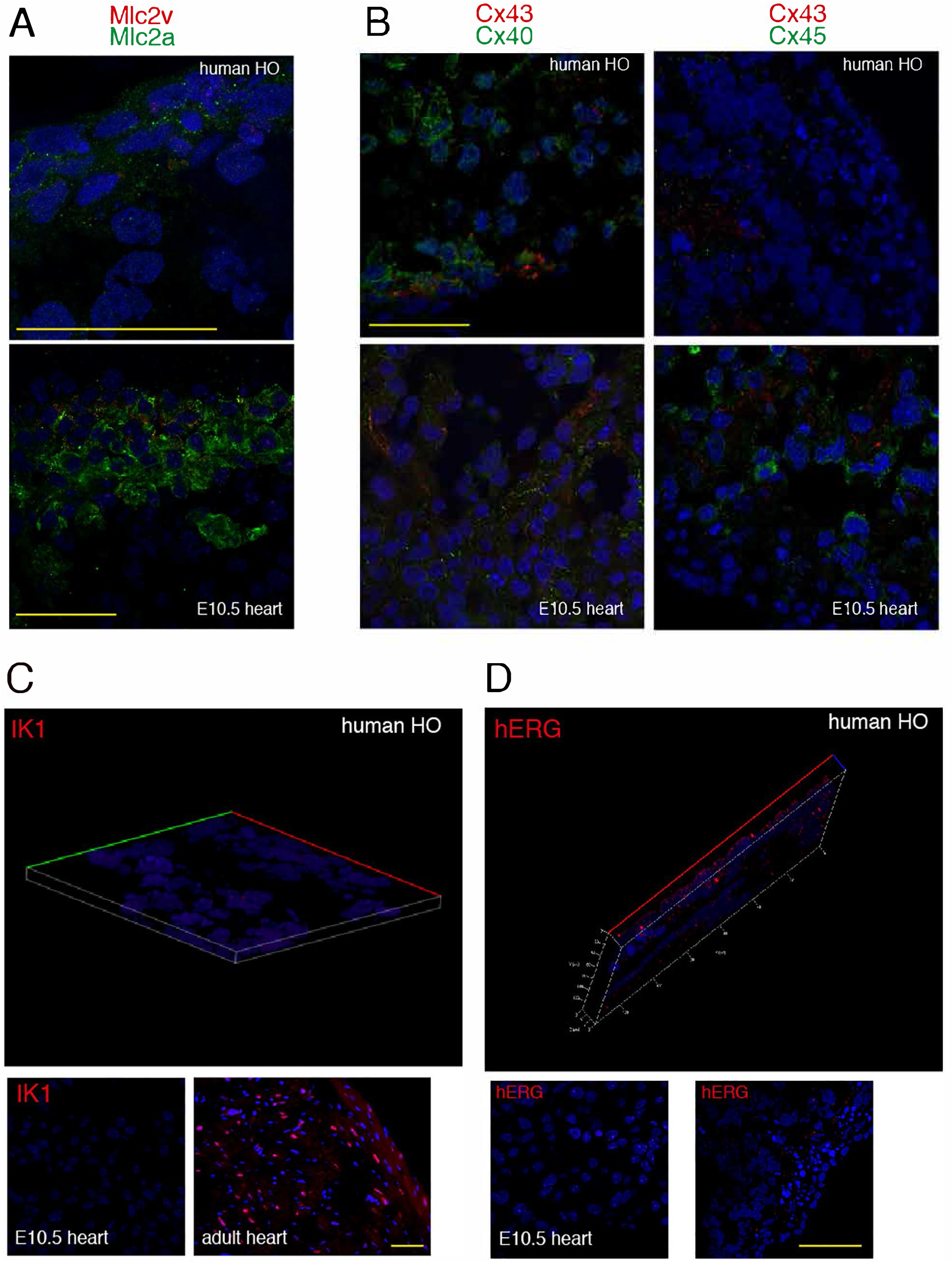
Evaluation of hESC-derived HOs. (**A**) Immunofluorescence staining for Mlc2v (red) and Mlc2a (green) in the hESC-derived HOs. Top: a representative hESC-HO. Bottom: mouse embryonic heart from E10.5. Scale bar: 50 μm. (**B**) Immunostaining to detect Cx43 (red)/Cx40 (green), (left panels), and Cx43 (red)/Cx45 (green) (right panels). Top: hESC-HOs. Bottom: mouse embryonic hearts from E10.5. Scale bar: 50 μm. (**C**) Immunostaining to detect one of potassium channel proteins, IK1 in a representative HO (top), E10.5 mouse embryonic heart (bottom left) and adult mouse heart (right, bottom). Scale bar: 50 μm. (**D**) hERG immunostaining in a representative HO (top and right, bottom), E10.5 mouse embryonic heart (left, bottom). Scale bar: 50 μm.

To identify the ultrastructural features of the hESC-derived heart organoids, we performed transmission electron microscopy (TEM) analysis. The intercalated disk (ID) is a cardiac muscle-specific ultrastructure required for coordinated muscle contraction and conduction of electrical excitability in the function of hearts (*8*). Additionally, the ID is composed of gap junctions, desmosomes, and facia adherens (intermediate junction) and arranged by a simple network or complex network (*17*). The hESC-derived heart organoids showed cardiac ultrastructural properties, including large mitochondria and abundant desmosomes around cardiac muscle filaments, glycogens, and secretory vesicles in atrial CMs. Notably, transverse sections of the hESC-derived heart organoids exhibited a complicated arrangement of IDs, suggesting that these human heart organoids have functional properties in terms of cardiac muscle contraction and conduction (fig. S3A).

To quantify cardiac contraction in hESC-derived heart organoids, we used MUSCLEMOTION software (*18*) with the images obtained by high-speed camera. The regular contractility was observed in hESC-derived heart organoids (fig. S3B), although the frequency of contraction was lower (6.8 times/20 sec, SEM = 0.77, n = 5) than that in mES-derived heart organoids (22.6 times/20 sec, SEM = 1.1, n = 4), suggesting the physiological discrepancy between human and mouse hearts.

For proper conduction, gap junction proteins such as Connexin 43 (Cx43, GJA1), Cx40 (GJA5) and Cx45 (GJC1) play essential roles in impulse propagation in the heart (*19*). In the human heart, Cx43 is expressed in atrial and ventricular working myocardial cells, Cx40 is mainly expressed in the atria, and Cx45 exists in His-bundle and bundle branches. To confirm the expression of gap junction proteins at the histological level, we performed immunofluorescence staining with the generated hESC-derived heart organoids. As expected, differential localization of Cx43, Cx40 and Cx45 was detected in the hESC-derived heart organoids, and the expression pattern of these gap junction proteins was similar to that of the mouse embryonic heart (E10.5) (Fig. 2B), suggesting adequate recapitulation of cardiac gap junctions in the hESC-derived heart organoids.

Because four types of ion channels containing K_ir_2.1, hERG, Na_v_1.5 and Ca_v_1.2 are especially important for the myocardial function as well as the development and the maturation of myocardial tissue in the heart, it is notable to determine whether these major ion channels are expressed in hESC-derived heart organoids. Inwardly rectifying potassium (K) channel K_ir_2.1, is predominantly expressed in the heart, and its expression during heart development gradually increases in the late embryonic and postnatal hearts. Importantly, K_ir_2.1 confers membrane stability and action potential repolarization in the cardiac cells, and produces the inward rectifier potassium current, I_K1_ current. Also, mutations of the gene encoding human K_ir_2.1 (KCNJ2) are responsible for cardiac arrhythmia in Andersen’s syndrome (*20*),(*21*). Therefore, we examined the expression of K_ir_2.1 in the hESC-derived heart organoids to determine whether these heart organoids indicate appropriate electrophysiological features like developing hearts *in vivo*. Immunofluorescence staining showed strong and broad expression of K_ir_2.1 in all hESC-derived heart organoids examined (Fig. 2C), suggesting that the resting membrane potential and cardiac excitability of individual cardiomyocytes of these heart organoids might be controlled by I_K1_.

Additionally, mutations of the human ether-a-go-go-related gene (HERG), which encodes the major subunit for the hERG channel, also cause cardiac arrhythmia as well as mutations of KCNJ2 (*22*). Given that hERG as one of voltage gated K channels produces I_Kr_ which is the rapidly activating component of I_K_, the major outward current involved in the repolarization of action potential in human heart muscle cells (*23*), we investigated the expression of hERG channel in the hESC-derived heart organoids. Immunofluorescence staining exhibited clear expression of hERG channels in all hESC-derived heart organoids examined (Fig. 2D), suggesting that the hESC-derived heart organoids generated in this study might produce functional I_Kr_ via hERG channel.

Regarding the development of new therapeutic drugs, the investigation of hERG channels’ block which causes a harmful side effect in human hearts is important for the preclinical test of drugs for their safety. For example, pentamidine, an antiprotozoal compound treated in Pneumocystis pneumonia causes acquired long QT syndrome by the direct block of I_Kr_ current. This block of I_Kr_ current occurred by the inhibition of hERG channels export from endoplasmic reticulum (ER) by pentamidine (*24*). Therefore, we examined whether hESC-derived heart organoids exhibit the inhibition of hERG channel trafficking in cell surface by the treatment of pentamidine. Thirty mM pentamidine were added in 17 days cultured hESC-derived heart organoids for over 16 hours (overnight). To detect the localization of hERG in the cells of heart organoids, we immunostained hESC-derived heart organoids with a hERG antibody and an ER marker, KDEL antibody. Expectedly, the export of immature hERG from ER was inhibited in hESC-derived heart organoids incubated with pentamidine, while the strong hERG signals were detected in the outside of ER in hESC-derived heart organoids without pentamidine (fig. S4), suggesting the faithful reflection of drug response on hERG channel trafficking block in hESC-derived heart organoids.

In normal cardiac functions, not only the outward currents but also the inward currents like the Na current and the Ca current are important. Among them, the inward Na current, which initiates action potentials in the heart, plays a crucial role in fast impulse propagation in cardiac tissue.

The main cardiac sodium channel is Na_v_1.5, which is encoded by SCN5A (*25*). Mutations in SCN5A are associated with multiple heart diseases, such as Brugada syndrome, long QT syndrome, conduction disease and cardiomyopathy. Additionally, genetic variation in SCN5A is associated with differences in cardiac conduction and the risk of arrhythmia. To determine the expression of Na_v_1.5 in the hESC-derived heart organoids, we performed immunofluorescence staining with an anti-SCN5A antibody. We detected clear expression of SCN5A in the hESC-derived heart organoids, suggesting the fast depolarization phase of cardiac action potential (Fig. 3A).

**Fig. 3.**
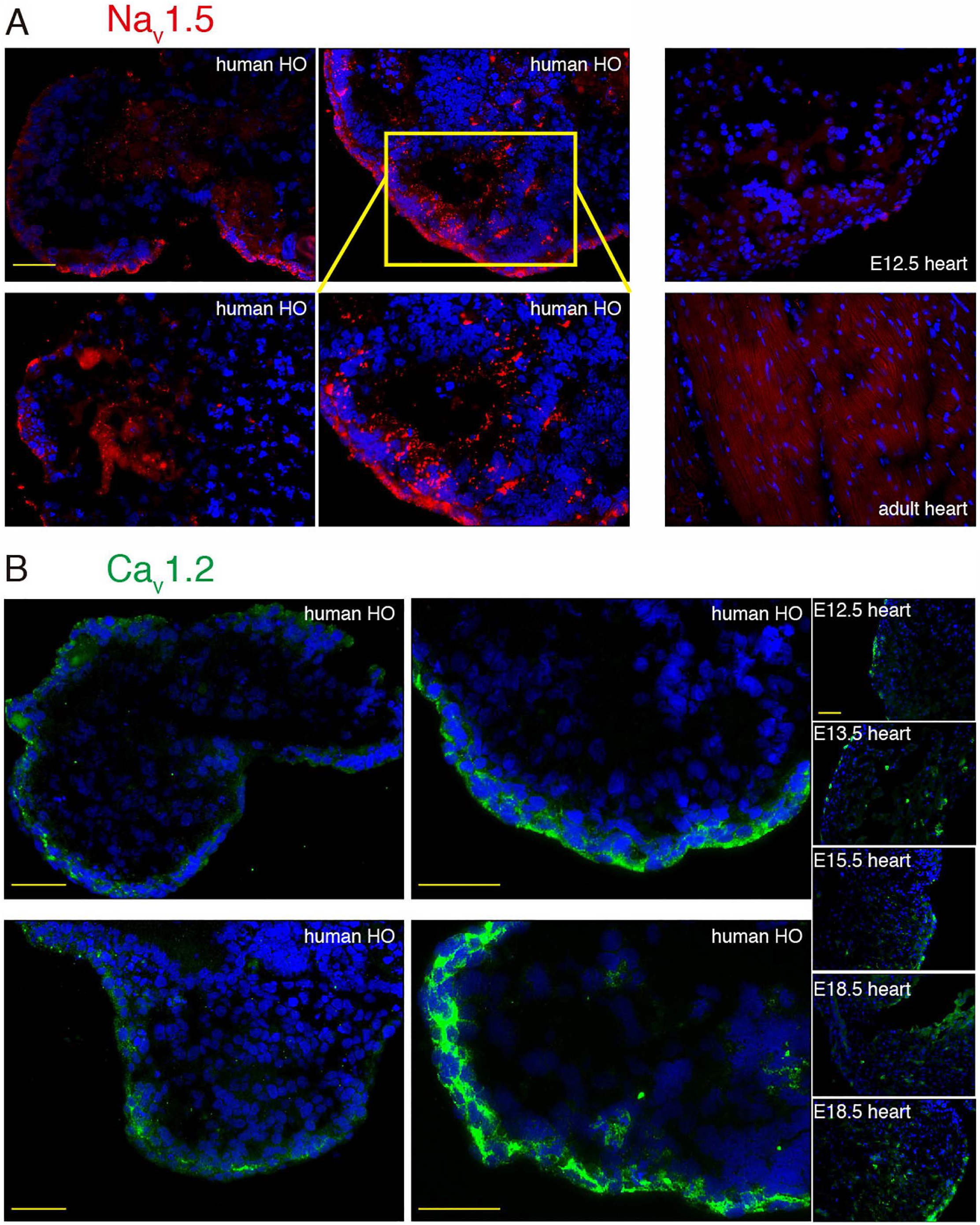
Expression of sodium and calcium channel proteins (Na_v_1.5 and Ca_v_1.2) in hESC-derived HOs. (**A**) Na_v_1.5 immunostaining to detect sodium channel in hESC-derived HOs (left and center), E10.5 mouse embryonic heart (top right), mouse adult heart (bottom right). Scale bar: 50 μm. (**B**) Ca_v_1.2 immunostaining to detect calcium channel in hESC-derived HOs (left and center), E12.5, E13.5, E15.5 and E18.5 mouse embryonic heart (right). Scale bar: 50 μm.

Another major inward current is Ca current. In cardiovascular physiology, voltage gated calcium channels including Ca_v_1.2 and Ca_v_1.3 (L-type Ca_v_channel) play important roles for calcium influx by Ca_v_1.2 in adult cardiomyocytes and contribute to inward current during cardiac action potential and excitation-contraction coupling (*26*). Ca_v_1.2 which is encoded by CACNA1C expresses in ventricular and atrial cardiomyocytes. Mutations of CACNA1C are associated with long-QT syndrome (gain of function) and Brugada syndrome (loss of function). Given the significant contribution of Ca channel to normal cardiac function, we next examined the expression of Ca_v_1.2 in hESC-derived heart organoids. Immunofluorescence staining with an anti-CACNA1C antibody showed strong expression of CACNA1C in hESC-derived heart organoids (Fig. 3B), suggesting the cardiac contractility triggered by Ca influx via Ca channels.

### Physiological ion currents in the hESC-derived heart organoids

Recent studies showed the importance of normal Na_v_1.5 channel activity for not only cardiac excitability but also histological development (*27*), therefore, the identification of functional Na_v_1.5 ion current is thought to be essential to ensure the normal cardiac development of hESC-derived heart organoids. To characterize the electrophysiological properties of ion current, patch clamp methods are golden standard and most reliable, especially, for the isolation of specific ion current from multiple ion currents in stem cell derived cardiomyoctes. Therefore, to analyze the electrophysiological features of the CMs prepared from the hESC-derived heart organoids cultured 19 to 36 days after EB formation, we enzymatically isolated the CMs and performed whole-cell voltage-clamp experiments with the isolated CMs. In many cells from heart organoids, typical inward currents, which had the maximal peak current at -20 mV (n=4), were induced in response to depolarizing step pulses (< 100 pA: n = 6; > 100 pA, < 500 pA: n= 13; > 500 pA: n = 8; n = 26 in total), suggesting that the induced inward currents were mainly mediated by voltage-dependent Na channels (Fig. 4A). These results suggested proper recapitulation of impulse propagation in the heart organoids by action potential initiation via Na currents. The inward currents were substantially reduced by changing to a Na-free extracellular solution (Fig. 4B top) or by inclusion of tetrodotoxin (TTX), a specific inhibitor of Na channels (Fig. 4B bottom). Ca currents were barely but clearly observed in a few cells from the hESC-derived heart organoids, each of which had an amplitude of approximately 40 pA following the blockade of Na currents (Fig. 4B top and bottom). The remaining inward currents showed even slower activation kinetics, which are characteristic of voltage-dependent Ca channels. Further reduction in the remaining inward currents was achieved by cobalt, an inorganic Ca channel blocker. Thus, the isolated cells from the hESC-derived heart organoids express ion channels that play integral roles in shaping cardiac action potentials; that is, voltage-dependent Na and Ca channels contributing to the initial rapid depolarization (phase 0) and plateau phase (phase 2) in cardiac action potentials, respectively.

**Fig. 4.**
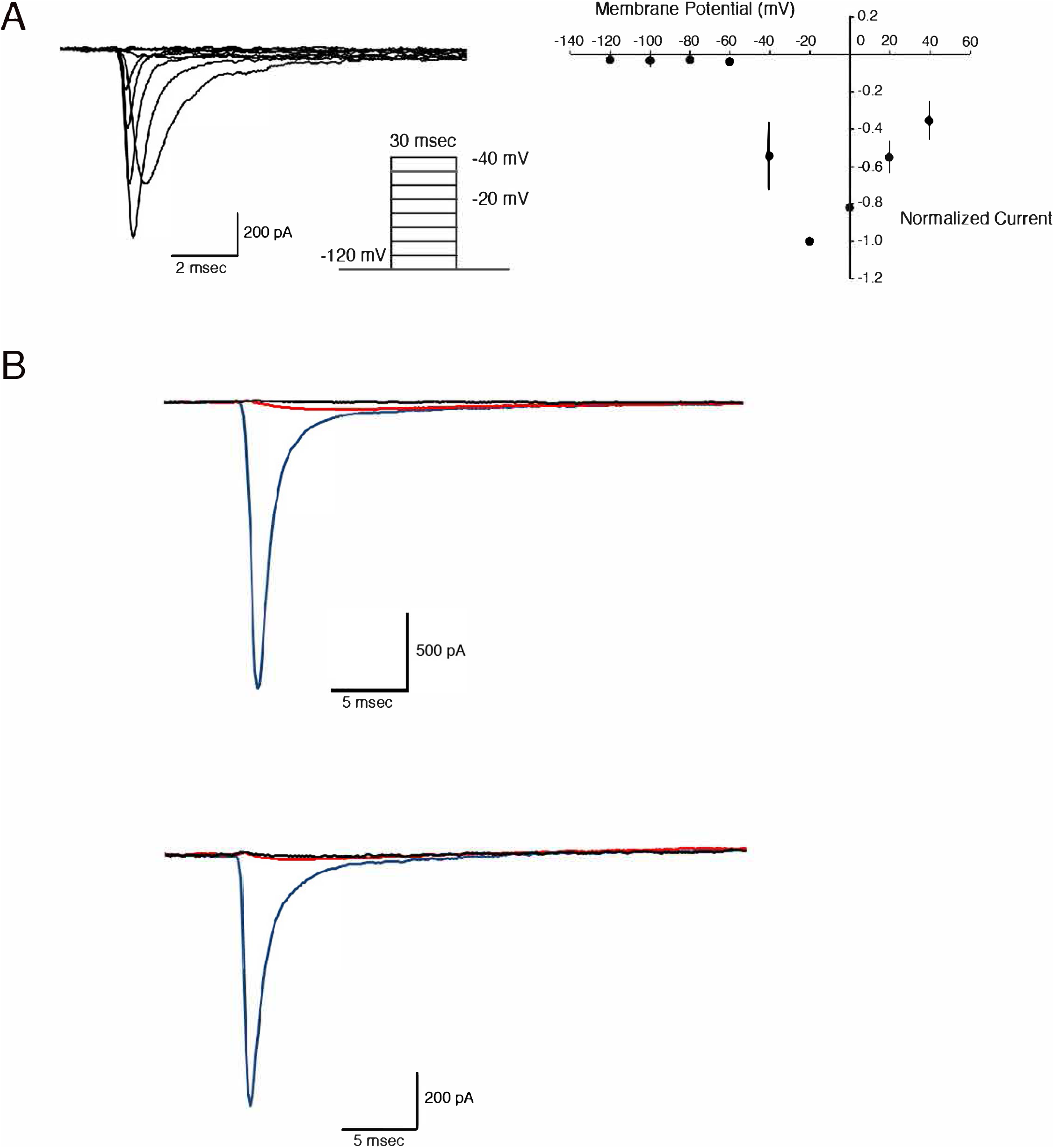
Electrophysiological analyses of CMs isolated from hESC-derived HOs in the whole-cell patch-clamp. (**A**, left) Representative Na current traces were elicited by the depolarizing voltage protocol, as shown in the inset. (**A**, right) Current-voltage relationships of peak I_Na_ with maximal inward currents at -20 mV (n = 4). Normalized peak amplitudes of I_Na_ to the maximal value were plotted as a function of the membrane potential. (**B**) Inward currents (blue) were substantially reduced by replacing Na ions with NMDG (top, red) or inclusion of TTX at 20 μM (bottom, red), and the addition of CoCl at 4 mM abolished the residual currents (top and bottom, black).

## Discussion

Heart development starts at a very early stage of embryogenesis (cardiac crescent at the E7.5 heart in mice corresponding to 20 days of gestation in humans) with a limited cell number and simple extracellular environment and proceeds to distinctive morphogenesis with functionality. Thus, elucidation of the spatiotemporal regulation of cardiogenesis is important for understanding not only heart development but also the structure-related specific functions of each part of the developing heart as well as the complete regulatory system of organogenesis in the body. In our previous study, we successfully generated mouse ESC-derived 3D heart organoids comprised of CMs, ECs and SMCs by inducing self-organization of cells in the presence of exogenous growth signals and extracellular matrix (*8*). The derived structures exhibited morphological changes such as chamber formation similar to E10.5 heart in mice corresponding to 29 days of gestation in humans, involving multiple structural heart cell types in a developmental stage-specific manner and showed cardiac ultrastructural properties and autonomous beating with myocardial contraction, suggesting the acquisition of functionality. As a biomimetic model, organized heart organoids can be promising tools for understanding cardiogenesis in many species.

The demand for human heart organoids in drug discovery as well as cardiac etiology and pathology has increased. Therefore, we applied the self-organized heart organoid-generating method to human ESCs with slight modifications and obtained hESC-derived heart organoids with multiple chambers and the expression of heart-specific proteins, including gap junction proteins and transcendent electrophysiological properties. Our results clearly indicated that FGF4 and LN/ET extracellular matrix trigger early cardiac development in humans and mice, suggesting that these two factors have conserved roles among mammalian species in adequate cardiac morphogenesis.

Importantly, the Na and Ca ion currents were detected in CMs of the hESC-derived heart organoids through patch clamp analysis, suggesting *in vitro* reconstitution of the human heart, along with action potential propagation and cardiac contraction. In particular, the Na ion channel protein, Na_v_ 1.5, was detected in hESC-derived heart organoids due to the normal Na ion current. These findings might lead to the development of a system for Na ion current blockade using hESC-derived heart organoids for drug toxicity screening; nevertheless, further investigations of hESC-derived heart organoid cells are required to determine their functional maturity, such as differentiation into cardiac cell types, developmental stages, and excitation-contraction (EC) coupling.

Although heart beating of the hESC-derived heart organoids was weaker than mouse heart organoids, MUSCLEMOTION analysis demonstrated they surely exhibit regular repeated contractile and relaxant responses, thus, suggesting that they are a promising tool for understanding cardiogenesis as well as *in vitro* drug safety test system.

The hESC-derived heart organoids have a remarkable cardiac morphology with atrial-like and ventricular-like structures that reflects adequate function, suggesting dramatic morphogenic changes in cardiogenesis by beating CMs or contraction that induce morphogenic shifts by mechanical force stress load. This issue might be solved by a mechanical stress culturing system that accelerates (stimulates) mechanical force and elongation stretch. Lipid metabolism is also important in heart function and the maturation of CMs. During mouse embryonic development, glycogens are abundant in embryonic CMs and are supplied as energy sources. However, cAMPs are generated by lipid metabolism in mature CMs, showing normal cardiac performance during the postnatal period. Additionally, palmitates were shown to be involved in the maturation of CMs induced from human pluripotent stem cells (*28*). Thus, further studies will be necessary to improve the maturity of CMs in hESC-derived heart organoids by switching the energy source from glucose to lipids depending on the developmental stage and applying mechanical loads in the culture system.

The heart is an essential organ for life maintenance together with the blood and circulatory system to carry nutrients and waste around the body to maintain its cells. Many patients suffer from heart diseases and defects, and the adult human heart cannot completely recover from damage. From another medical point of view, cardiovascular toxicity is one of the major barriers to drug development: unanticipated heart toxicity has been a major reason for the withdrawal of many anticancer drugs and anti-inflammatory drugs from the market. Therefore, a new and safe medical treatment for many diseases and an effective nonclinical heart toxicity evaluation system closer to the *in vivo* heart are eagerly anticipated. Thus, pluripotent stem cell-derived heart organoids will be indispensable for nonclinical heart toxicity evaluation systems, including both patch clamp analysis from isolated cells and drug responses of the heart organoids as a whole.

Finally, a deep understanding of the heart regulatory system using hESC-derived heart organoids will provide important information on not only the etiology of cardiovascular diseases, thereby leading to the development of a new therapeutic method, but also the developmental relationships among organs in our body.

## Acknowledgments

The authors would like to thank Drs. T. Furukawa (TMDU), T. Sasano (TMDU), and K. Ihara (TMDU) for helpful discussions. The authors would like to thank Dr. H. Akutsu (National Center for Child Health and Development) for providing the cell lines of hESCs.

## Author contributions

Conceptualization: JL, FI

Methodology: JL, HM, KS

Investigation: JL, HM, RK

Visualization: JL, HM

Funding acquisition: JL

Project administration: JL, FI

Supervision: JL, KS, FI

Writing – original draft: JL, HM

Writing – review & editing: JL, HM, KS, FI

## Supplementary Materials

Materials and Methods

Figs. S1 to S4

References (*29*–*31*)

Movies S1 to S2

