## Supplementary Materials for "*In vitro* generation of human embryonic stem cell-derived heart organoids possessing physiological ion currents"

**This PDF file includes:**

Materials and Methods
Figs. S1 to S4
Captions for Movies S1 to S2

**Other Supplementary Materials for this manuscript include the following:**

Movies S1 to S2

#### Materials and Methods

Animals and human embryonic stem cells (hESCs)

All animal experiments were approved by the Institutional Animal Care and Use Committee of Tokyo Medical and Dental University (TMDU) and were performed according to the guidelines of this institution. All experiments using hESCs were approved by the institutional hESC Use Committee of TMDU and were performed according to the guidelines of this institution.

Cell culture

The hESCs (gifted from Dr. Akutsu, National Research Institute for Child Health and Development, Japan) were cultured and maintained with Primate ES cell medium (Reprocell) supplemented with bFGF (final concentration 10 ng/mL) on SL10 feeder cells (Reprocell). Prior to EB formation, the hESCs were cultured twice on Matrigel (0.3 mg/mL per 1-well of a 6-well plate, BD) to remove feeder cells. For hESC-derived EB formation, the cultured hESCs on a Matrigel coat, washed with PBS (++) and incubated with 0.5 mL of CTK solution (Reprocell) for 3 min at 37°C. The dissociated hESCs were collected by a cell scraper and seeded in a U-bottomed 96-well plate (Prime surface 96-well U, Sumiron) in KSR- and NEAA-supplemented DMEM Glutamax medium (KSR-EB medium) with Y27632 (10 mM) and incubated at 37°C for 1 day. On culturing day 1, the medium was changed to fresh KSR-EB medium without Y27632 and incubated until culturing day 4. For the generation of hESC-derived heart organoids, we performed the *in vitro* culture of hESC-derived EBs by using a protocol previously described (*8*). Briefly, hESC-derived EBs were placed onto 85.7 μg/cm$^2$ LN/ET gel (BD 354259)-coated chamber slides (Falcon 354118) in 200 μL of heart organoid basic medium [50 μg/mL penicillin/streptomycin, 20% KSR, 1 mM sodium pyruvate, 100 μM β-mercaptoethanol, 2 mM L-glutamine, 60 ng/mL progesterone, 30 ng/mL β-oestradiol, 5 μg/mL insulin, 20 μg/mL transferrin, and 30 nM selenite in DMEM/F12 (Gibco 11320033)] containing FGF4 (60 ng/mL). Subsequently, the hESC-derived EBs were incubated at 37°C and 5% $CO_2$ for 15 days or further with medium changes on days 3, 5, 7, 9, 11, 13, and 15. Starting on day 9, BMP4 (50 ng/mL), BIO (2.5 μM), LIF (1,000 units/mL) (*7*) (BBL), BIO, BMP4, BMP10, and LIF (BBBL) and additional supplementation with the ROCK inhibitor Y27632 (Y) were added to FGF4 heart organoid basic medium. Additionally, the hESC-derived heart organoids were transferred onto 10.7 μg/cm$^2$ iMatrix-411 (Laminin 411, Nippi)-coated chamber slides at culturing day 11. After culturing, the hESC-derived heart organoids were collected for further analyses, including immunofluorescence staining, ultrastructural analysis and patch clamp analysis.

Immunofluorescence staining

Immunofluorescence staining was performed as reported previously (*8*). hESC-derived heart organoids were collected after 15 days of culture and frozen in optimal cutting temperature (OCT) compound (Tissue-Tek) in a plastic tissue mould (Cryo dish, Shoei). The sections were sliced at 5 μm using a cryostat set at -16°C and transferred to MAS-coated glass slides (Matsunami). Additionally, mouse embryonic, postnatal and adult heart sections were prepared from C57BL/6 mouse embryos from 9.5 to 13.5 days postcoitum and postnatal day 1 (P1) and adults (over 9 weeks). Prior to freezing in OCT, the mouse hearts were immersed in sucrose-PBS solutions (4%, 10%, 15%, and 20%). The cryosections were fixed in 4% paraformaldehyde-PBS at room temperature (RT) for 15 min. The fixed sections were washed three times in PBS for 5 min and incubated with a blocking buffer (PBS containing 5% normal goat serum and 0.3%

Triton X-100 or PBS containing 5% bovine serum albumin (BSA) and 0.3% Triton X-100) at RT for 1 h. The sections were subsequently incubated overnight with primary antibodies in dilution buffer (PBS containing 1% BSA and 0.3% Triton X-100) at 4°C. After the sections were washed with PBS, they were incubated with secondary antibodies at RT for 1 to 2 h. The slides were counterstained with DAPI (diluted 1:1,000, Dojindo Laboratories) and mounted using a VECTASHIELD HardSet Antifade Kit (VECTOR Laboratories). Primary antibodies against the following proteins were used: Mlc2a (Synaptic System #311 011), Mlc2v (Synaptic System #310 003), Cx43 (Sigma, C6219), Cx40 (Invitrogen, #37-8900), Cx45 (ab78408), KCNN4 (IK1, $K_{ir}2.1$, GTX54786), and potassium channel $hK_v11.1$ (hERG, Sigma Aldrich, #P9497). The appropriate secondary antibodies (all from Invitrogen) for each primary antibody were also used.

Whole-mount immunostaining by the tissue clearing method

For whole-mount immunostaining by tissue clearing, we used the tissue clearing reagent CUBIC-L (TCI) as described in the manufacturer's protocol. The collected hESC-derived heart organoids were cultured for over 20 days, and mouse embryonic hearts were fixed in 4% PFA/PBS overnight at 4°C and washed with PBS three times for 2 h each at RT. The washed samples were delipidated by incubating in 750 mL of CUBIC-L solution in a shaker for 2 days at 37°C and then washed with PBS three times for 2 h each. For immunostaining, the samples were incubated in mixed antibodies (Alexa Fluor 647 Mouse Anti-cTnT; BD Phar, BV480 Rat Anti-Mouse CD31; BD Horizon, anti- aSMA antibody Alexa Fluor 568; Abcam) diluted with staining buffer (0.01% sodium azide and 0.5% Triton X-100 in PBS) for 3 days at RT and washed with PBS three times for 2 h each. For RI matching, the samples were incubated in 750 mL of CUBIC-$R^+$ solution overnight at RT. The immunostained hESC-derived heart organoids were analysed with a laser confocal microscope (Lightsheet, Zeiss).

HERG channel trafficking inhibition assay

For the inhibition assay of hERG channel trafficking, 17 days cultured hESC-derived heart organoids were cultured for over 16 hours (overnight) under conditions in the absence or in the presence of 30 mM of pentamidine prepared in ddw. After incubation with/without pentamidine, hESC-derived heart organoids were collected, embedded in OCT compound, and cryosectioned at 5 mm using a cryostat. The cryosections were used for immunostaining with anti-mouse KDEL (10C3) antibody (Novusbio, #NBP1-97469SS) and hERG antibody (Sigma Aldrich, #P9497) and appropriate secondary antibodies. The cryosection slides were counterstained with DAPI, mounted and observed the subcellular localization of hERG channels under laser confocal microscope (Zeiss, LSM710).

Electrophysiology

**Cell preparation**

Cells were prepared from hESC-derived heart organoids 18–30 days post-EB formation with an enzymatic digestion method as described previously (*29*). Briefly, hESC-derived heart organoids were washed with cell isolation buffer (CIB) twice, 200 µL of enzyme mix [collagenase II (1 mg/mL, Worthington), collagenase IV (1 mg/mL, Sigma), 0.3 mM $CsCl_2$ in CIB] was added, and the cells were incubated at 37°C for 15 min. The samples were mixed by pipetting (10 times), 500 µL of CIB-$Ca^{2+}$-BSA was added, and small pieces of samples were broken up into single cells by pipetting 7 times. The dissociated cells were centrifuged at 2000 rpm for 2 min, washed with CIB-$Ca^{2+}$-BSA, and finally centrifuged at 2000 rpm for another 2 min. The cell pellets were suspended in 100 µL of normal Tyrode solution (Sigma T2397). For attachment, isolated cells were transferred on round cover slips precoated with collagen type I (IWAKI, Tokyo, Japan) and maintained in Tyrode solution in a humidified culture dish for over 30 min until use for electrophysiological experiments.

**Patch-clamp**

A conventional patch clamp method was used in the whole-cell voltage-clamp mode(*30*). Cover slips with the isolated cells from hESC-derived heart organoids were transferred to a temperature-controllable recording chamber and superfused with the extracellular solutions at a rate of 2–3 mL/min. Pipette electrodes were fabricated from borosilicate glass capillaries by a filament using a P-97 micropipette puller (Sutter Instrument, Novato, CA, USA). The pipette electrodes with electrical resistance between 2 and 5 MOhm were used. Membrane currents mediated by ion channels were filtered at 2 or 5 kHz using an Axopatch 200B amplifier and digitized at 20 kHz using pCLAMP 10 Software Suite (Molecular Devices, San Jose, CA, USA). Whole-cell voltage-clamp configuration was obtained by visual access under an upright microscope (Olympus, Tokyo, Japan). All the recordings were performed at 30–33°C. The intracellular solution contained the following (in mM): 50 CsCl, 60 CsF, 10 NaCl, 2 $MgCl_2$, 20 ethylene glycol-bis (2-aminoethylether)-N,N,N',N'-tetraacetic acid (EGTA), and 10 4-(2-hydroxyethyl)-1-piperazine-ethanesulfonic acid (HEPES); the pH was adjusted to 7.25 with 1 M CsOH. The extracellular solution contained the following (in mM): 140 NaCl, 4 CsCl, 1 or 2 $CaCl_2$, 1.2 $MgCl_2$, 10 glucose, 10 HEPES; the pH was adjusted to 7.40 with 1 M NaOH. When Ca currents were recorded, another intracellular solution was used and contained the following (in mM): 112 CsCl, 27 CsF, 2 NaCl, 8.2 EGTA, 4 MgATP, 10 HEPES; the pH was adjusted to 7.25 with 1 M CsOH, and tetrodotoxin (TTX) was additionally included at 20 µM externally or a different extracellular composition was used as follows (in mM): 130 N-methyl-D-glucamine (NMDG), 4 CsCl, 4 $BaCl_2$, 1.2 $MgCl_2$, 10 glucose, 10 HEPES; the pH was adjusted to 7.40 with 1 M HCl. To ensure that the remaining currents after removing $Na^{\pm}$ currents were mediated by Ca channels, we added cobalt (Co) ions, which are known as an inorganic Ca channel blocker(*31*), to the extracellular solution at 4 mM. Organic and inorganic ion channel blockers, TTX and $CoCl_2$, were purchased from Fujifilm and Wako Pure Chemical.
For elicitation of inward currents through voltage-dependent ion channels, depolarizing step pulses were delivered at 0.1 Hz. Leak and capacitive currents were subtracted using a P/N protocol (8 times). P/N corrected waveforms were used to measure the peak amplitudes of the respective ion channels. Peak currents attained as incremental voltage pulses increased stepwise were normalized to the maximum value and plotted as a function of the voltages to generate current-voltage curves relationship (I-V curves). Peak analysis was performed using pCLAMP 10 Software Suite. Data are shown as the mean ± standard deviation.

### Figure S1

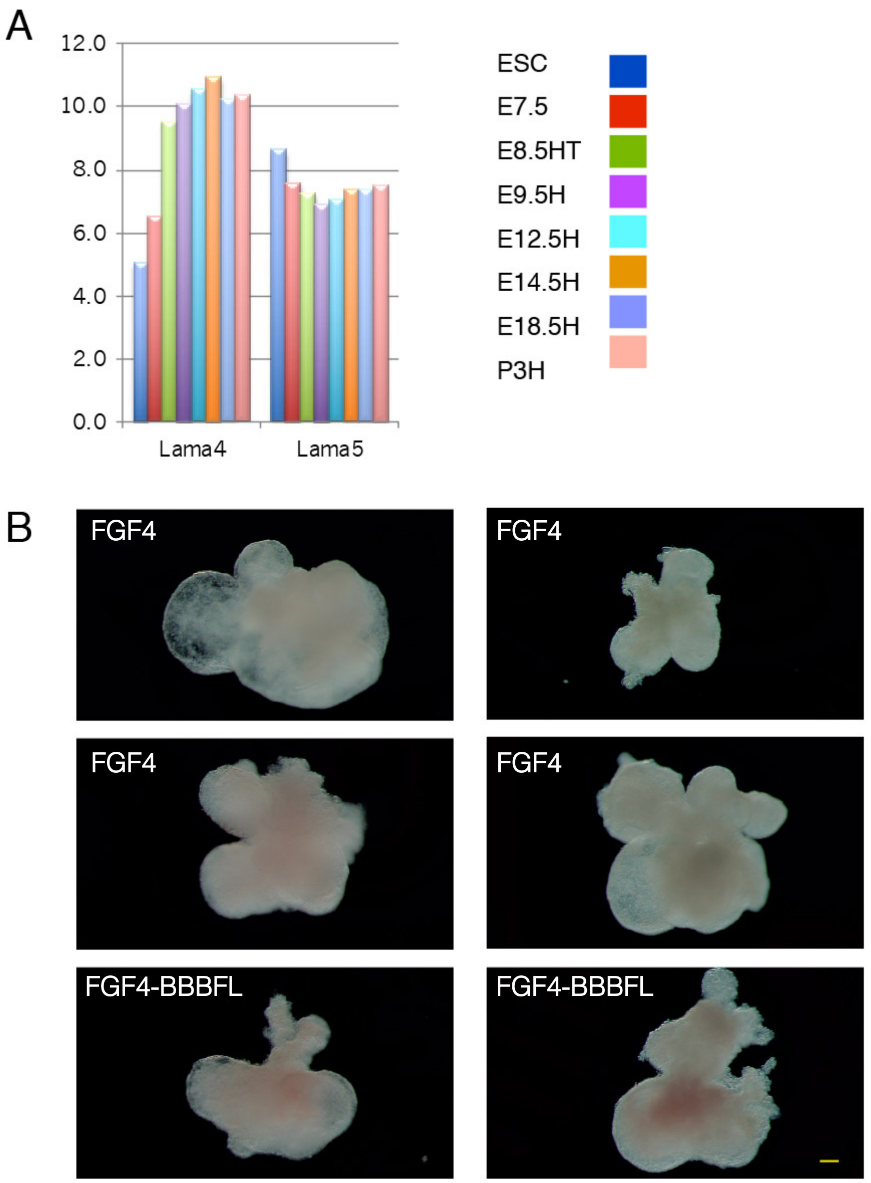


**Fig. S1.**
The requirements of hESC-derived HO generation. (A) Shift in the expression levels of the ECM genes Lama4 and Lama5 during mouse heart development. The GEO accession no. GDS5003 data 12 were reanalysed. to examine the expression changes in Lama4 and Lama5 in the mouse embryonic heart. (B) Successive chamber formation of hESC-derived HOs in the presence of FGF4 only and FGF4-BBBFL via self-organized morphogenesis. See also Fig. 1C. Scale bar: 100 μm.

Figure S2

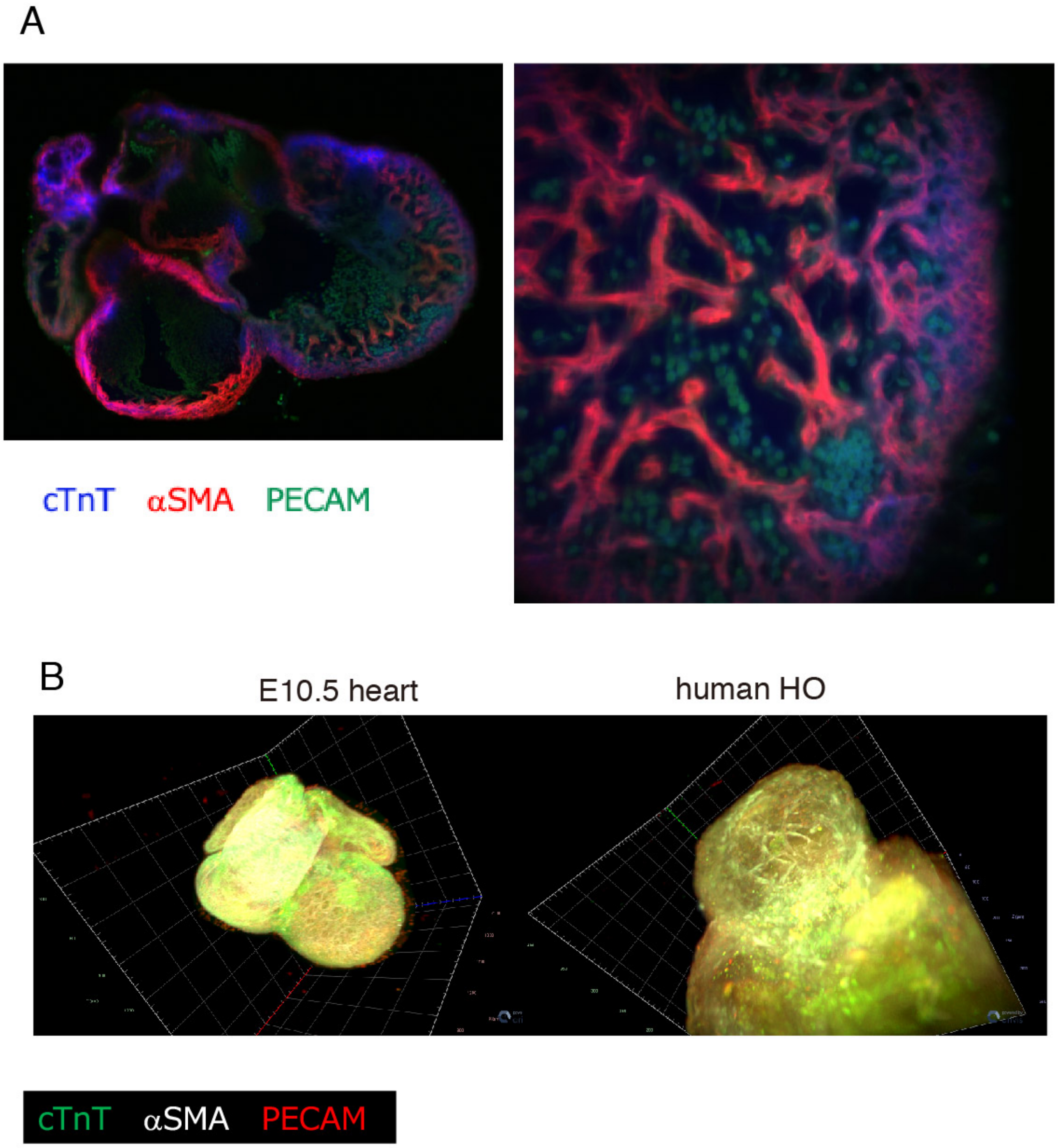


**Fig. S2.**
Differentiation of EBs into CMs, ECs, and SMs in hESC-derived HOs. (A) Wholemount immunofluorescence staining by the cubic tissue clearing method to detect PECAM (an EC marker, green), cTnT (a CM marker, blue), and an SM marker (red) in the embryonic heart from E10.5. Left: whole, right: magnified image of a ventricle. (B) Whole-mount immunofluorescence staining by the cubic tissue clearing method to detect PECAM (red), cTnT (green), and αSMA (white). Segregated cTnT expression (green) and/or broad expression of cTnT in myofibrils and its adjacent expression of PECAM (red) and αSMA (white) were detected in the hESC-derived HOs. These expression levels in both mouse E10.5 hearts (left) and hESC-HOs (right) exhibit distinctive reticulated structures of myofibrils and ECs.

### Figure S3

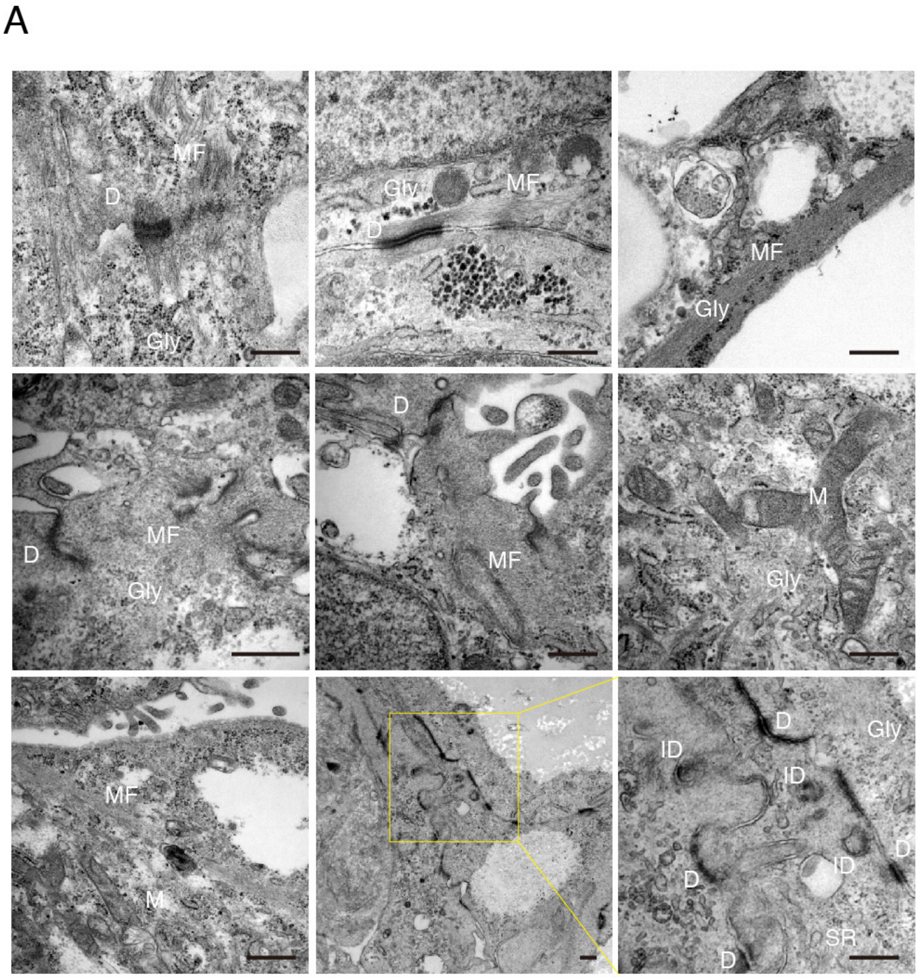


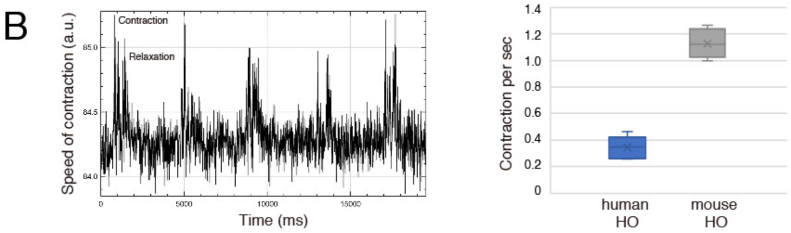


**Fig. S3.**

Ultrastructural features and cardiac contractility of hESC-derived HOs. (A) TEM analysis of the hESC-derived HOs shows CM-like ultrastructural properties, including islets of intercalated discs (ID), myofibril (MF), mitochondria (M), glycogens (Gly) in myofibrils, sarcoplasmic reticulum (SR) and desmosomes (D). Scale bar: 400 nm. (B) Contraction analysis of hESC-derived HOs by using MUSCLEMOTION software. The left histogram indicates a representative contraction and relaxation pattern in hESC-derived HOs and the right box plot indicates contraction per one second between human HOs and mouse HOs.

Figure S4

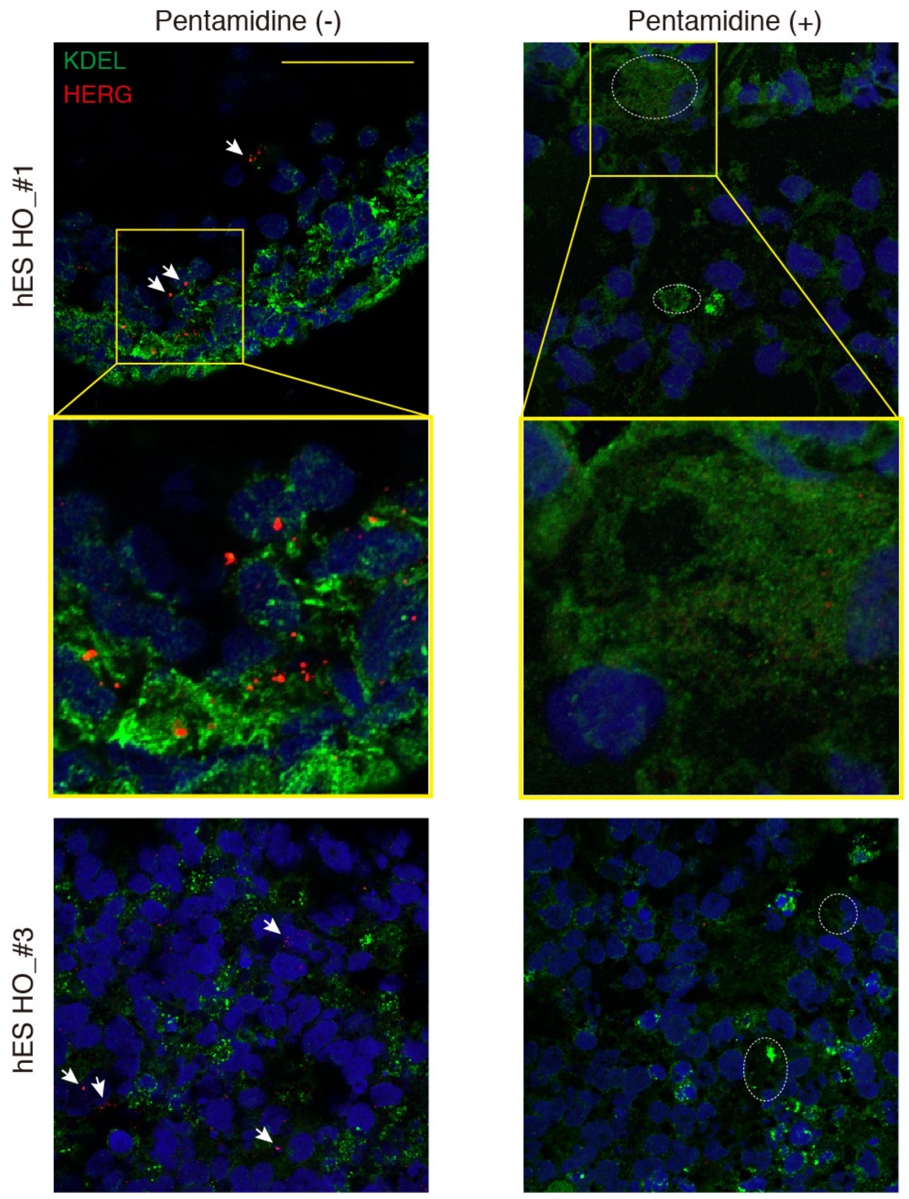


**Figure S4.**

HERG trafficking inhibition assay in hESC-derived heart organoids. The localization of hERG protein in hESC-derived heart organoids after the treatment with/without pentamidine at 30 μM were investigated. The sections of hESC-derived HOs were immunostained with anti-KDEL antibody used for detection luminal ER proteins and anti-hERG antibody. In the cells of hESC-derived heart organoids without treatment of pentamidine, hERG immunosignals were frequently localized outside ER and/or in the cell surface (arrow heads). In the cells of pentamidine treated hESCderived heart organoids, hERG immunosignals were mostly expressed in ER (dotted circle). Scale bar: 50 μm.

**Movie S1.**

Spontaneous beating of in vitro cultured heart organoid at day 11.
Human ESCs derived EB cultured under the condition of heart organoid generation protocol for 11 days. Multi-chambered heart organoid showed spontaneous beating movement indicating functional capacity of heart organoid. QuickTime movie.

**Movie S2.**

Spontaneous beating of in vitro cultured heart organoid at day 15.
Human ESCs derived EB cultured under the condition of heart organoid generation protocol for 15 days. Multi-chambered heart organoid showed spontaneous beating movement indicating functional capacity of heart organoid. QuickTime movie.
